# Identification of cells expressing Calcitonins A and B, PDF and ACP in *Locusta migratoria* using cross-reacting antisera and *in situ* hybridization

**DOI:** 10.1101/2021.07.28.454216

**Authors:** Jan A. Veenstra

**Affiliations:** INCIA UMR 5287 CNRS, Université de Bordeaux, Pessac, France

**Keywords:** Calcitonin A, Calcitonin B, ACP, PDF, affinity column, Locust

## Abstract

This work was initiated because an old publication suggested that electrocoagulation of four paraldehyde fuchsin positive cells in the brain of *Locusta migratoria* might produce a diuretic hormone, the identity of which remains unknown, since none of the antisera to the various putative *Locusta* diuretic hormones recognizes these cells. The paraldehyde fuchsin positive staining suggests a peptide with a disulfide bridge and the recently identified *Locusta* calcitonins have both a disulfide bridge and are structurally similar to calcitonin-like diuretic hormone.

*In situ* hybridization and antisera raised to calcitonin-A and -B were used to show were these peptides are expressed in *Locusta*. Calcitonin-A is produced by neurons and neuroendocrine cells that were previously shown to be immunoreactive to an antiserum to pigment dispersing factor (PDF). The apparent PDF-immunoreactivity in these neurons and neuroendocrine cells is due to crossreactivity with the calcitonin-A precursor. As confirmed by both an PDF-precursor specific antiserum and *in situ* hybridisation, those calcitonin-A expressing cells do not express PDF.

Calcitonin B is expressed by numerous enteroendocrine cells in the midgut as well as the midgut caeca. A guinea pig antiserum to calcitonin A seemed quite specific as it recognized only the calcitonin A expressing cells. However, rabbit antisera to calcitonin-A and-B both crossreacted with neuroendocrine cells in the brain that produce ACP, this is almost certainly due to the common C-terminal dipeptide SPamide that is shared between *Locusta* calcitonin-A, calcitonin-B and ACP.

## Introduction

Animals use a large diversity of hormones to regulate a variety of physiological processes. Many hormones are produced by specialized glands, such as the thyroid in vertebrates or the corpus allatum and prothoracic gland in insects. The existence of a gland making a single hormone allows for extirpation as well as its subsequent reimplantation, thereby factilitating studies into the physiological effects of its hormone(s). Other hormones are produced by specialized neuroendocrine cells within the nervous system. In those cases the specific elimination of the cells producing a hormone are more difficult to perform, although there are a few notable examples, such as the growth hormone producing cells in the pond snail *Lymnaea stagnalis* (Geraerts, 1976), those making prothoracicotropic hormone cells in the moth *Manduca sexta* (Agui et al, 1979), or the ones responsible for the production of virginoparae in aphids (Steel and Lees, 1977). Most cases where neuroendocrine cells were identified as the source of a specific hormone and thereby shown to be responsible for a particular function involved antisera to the hormone after its chemical structure had been established. Indeed, there are very few insect neuroendocrine cells for which a function has been established before the hormone itself had been identified. An interesting example concerns eclosion hormone in larvae of the dragonfly *Aeshna cyanea* by Charlet and Schaller (1976). This identification was indirectly confirmed several years later by the localization of *Manduca sexta* eclosion hormone in homologous neuroendocrine cells (Horodyski et al., 1989). Previously Charlet (1974) had shown that the electrocoagulation of another neuroendocrine cell group nearby led to an increase in body size in *Aeschna* larvae. A few years earlier a similar increase in body size had been reported after the elimination of homologous cells in *Locusta migratoria* (Girardie, 1970) and was subsequently also described for the cockroach *Rhyparobia maderae* (de Bessé, 1978), which was then still placed in the *Leucophaea* genus. The apparent increase in hemolymph volume and pressure led these authors to attribute this to the production and release of a diuretic hormone by those cells. Once these cells were eliminated, this would lead to an increase in hemolymph volume.

In insects primary urine is produced by the Malpighian tubules and water and ions are subsequently reabsorbed in the hindgut to produce the final urine (*e*.*g*. Coast et al., 2002). Because non peptide analogs might make powerful pesticides, a lot of research has been dedicated to the identification of insect diuretic hormones in insects. These hormones increase the rate of fluid secretion by the Malpighian tubules and a number of such peptides have been identified so far. The best known of these include the CRF-like diuretic hormone, DH31 (also known as the calcitonin-like diuretic hormone), leucokinin, tachykinin, periviscerokinins, also known as capa peptides, and an anti-parallel dimere of inotocin, a vasopressin-related peptide. With the exception of the inotocin dimere all of these peptides have been shown to increase fluid secretion by the Malpighian tubules in a variety of insect species (*e*.*g*. Proux et al., 1987; Kataoka et al., 1989; Lehmberg et al., 1991; Coast et al., 1993; Thompson et al., 1995; Johard et al., 2003; Davies et al., 2013; Furuya et al., 2000). Furthermore, in *Drosophila melanogaster* the Malpighian tubules express receptors for yet other neuropeptides and at least some of those peptides also stimulate fluid secretion (Chintapalli et al., 2012). Finally, 5-hydroxytryptamine is also able to stimulate fluid secretion by the Malpighian tubules, including in *Locusta* (Morgan and Mordue, 1984).

It remains unclear whether any of these peptides is an authentic diuretic hormone in the sense that it is used to eliminate excess water. Indeed, for some of these peptides this seems unlikely. Thus, whereas tachykinin stimulates the rate of fluid secretion by the Malpighian tubules in *Locusta migratoria* (Johard et al., 2003), it is during starvation that it is released into the hemolymph (Winther and Nässel, 2001), but it seems unlikely that starved locusts would have an excess of water. The periviscerokinins on the other hand stimulate fluid secretion by the Malpighian tubules in some species, but inhibit it in others. It has previously been suggested that the periviscerokinins together with the tryptopyrokinins and pyokinins are regulators of different aspects of digestion, which may benefit from increased hemolymph clearance associated with a higher activity of the Malpighian tubules (Veenstra et al., 2012).

Although antisera to the putative diuretic hormones mentioned above exist and have been tested on locusts and cockroaches, none of them have been described to label the neuroendocrine cells described above (*e*.*g*. Rémy and Girardie, 1980; Patel et al., 1994; Muren et al., 1995; Predel et al., 2007).

Girardie, Charlet and de Bessé all used a paraldehyde fuchsin staining method to identify those cells, which reacts with disulfide bridges to yield a blueish purple color. For many of the insect neurondocrine cells that were thus identified we now know which neuropeptides they produce. These include insulin, neuroparsin, bursicon, eclosion hormone, GPA2/GPB5, CCAP, inotocin, PTTH, and SIFamide. Most of these neuropeptides, or their precursors, contain several disulfide bridges, although in the case of SIFamide, there is only one and these cells stain only very weakly. It thus seemed likely that the precursor of the active neuropeptide produced by the cells identified by Girardie contained at least one disulfide bridge. Although the literature on insect brain neuroendocrine cells using paraldehyde fuchsin staining is very extensive, cells homologous to those in *Locusta* have never been described from holometabolous species.

The genome sequences of *L. migratoria*, the termite *Zootermopsis nevadensis* and the cockroach *Periplaneta americana* (Terrapon et al., 2014; Wang et al., 2014; Zeng et al., 2021) permit a much more complete analysis of their neuropeptides genes than was previously possible. One of the discoveries concerned the existence of an arthropod calcitonin gene that at least in the termite, the cockroach and decapod crustaceans, is alternatively spliced to produce what have been called the calcitonin A and calcitonin B neuropeptides (Veenstra, 2014, 2016; Zeng et al., 2021). In *L. migratoria*, however, these two types of calcitonin are produced from two different genes (Hou et al., 2015).

Transcriptome data from both decapods and phasmids suggested that calcitonin A is specific for the nervous system, while calcitonin B is abundantly expressed in the intestin (Veenstra, 2014, 2016). The presence of a disulfide bridge in calcitonin A, its absence from holometabola, its presence in the nervous system and its structural similarity to two putative insect diuretic hormones (DH31 and the CRF-like diuretic hormone) suggests this might be the peptide produced by the neuroendocrine cells identified by Girardie in 1970. Thus the identification of the cells producing calctionin seems of interest.

Preliminary results showed co-localization of calcitonin with pigment-dispersing factor (PDF) immunoreactivity, while antisera raised to calcitonins A and B recognized ACP. It is for this reason these two these peptides are also included in this study.

## Materials and Methods

### Insects

*Locusta migratoria, Schistocerca gregaria* and *Gryllus bimaculatus* were purchased at a local pet shop and from grillonshop.fr. Locusts were fed grass that was freshly cut on campus and oat brain, while crickets were supplied with water and mouse chow. *Periplaneta americana* came from my small laboratory colony and were maintained on mouse chow and water.

### Peptides

The Macromolecular Structure Facility at the University of Arizona produced NSEIINSLLGLPKVLNDAamide, *i*.*e. Gryllus* PDF, while I was employed there. This peptide was only used to test antiserum specificity. CAQGEDQHLNSIDSPamide was custom synthetized by Shanghai RoyoBiotech Co., Ltd (Shanghai, China) and TQYDEEKYQENEVC, CAERQQVASHDVamide, QAQGEDQHLNSIDSPamide, LTVDGARLRDRDA and GASDDGLYFESGSSPamide by PhtdPeptides Co., Ltd (Zhengzhou City, China).

### Antisera

Rabbit antisera to crustacean hyperglycemic hormone (CHH) that recognizes ITP in *Locusta* (Dircksen, 2009) and to decapod PDF (Dircksen et al., 1987) were donated by Heinrich Dircksen and a monoclonal antibody to *Drosophila* PDF (Cyran et al., 2005) was a gift from Justin Blau. Antiserum to *Locusta* CRF-like diuretic hormone (Patel et al., 1994) was a gift from Geoffrey Coast. A mouse polyclonal antiserum to ACP (Patel et al., 2014) was also used, as were rabbit antisera to corazonin (Veenstra and Davis, 1994), leucokinin (Chen et al., 1994), allatotropin (Veenstra and Hagedorn, 1993) and myoinhibitory peptide (Park et al., 2008).

Guinea pig antiserum was raised to the predicted PDF associated RYamide using TQYDEEKYQENEVC. Two calcitonin A antisera were produced, one in a guinea pig with QAQGEDQHLNSIDSPamide and the second one in a rabbit using CAQGEDQHLNSIDSPamide. Other rabbit antisera were produced against CAERQQVASHDVamide (HDVamide) and LTVDGARLRDRDA (RDRDA), the C-terminal sequences of two other peptides predicted to be produced from the calcitonin A precursor and GASDDGLYFESGSSPamide, the C-terminal sequence of *Locusta* Calcitonin B1 (Fig. 1).

**Figure 1.**
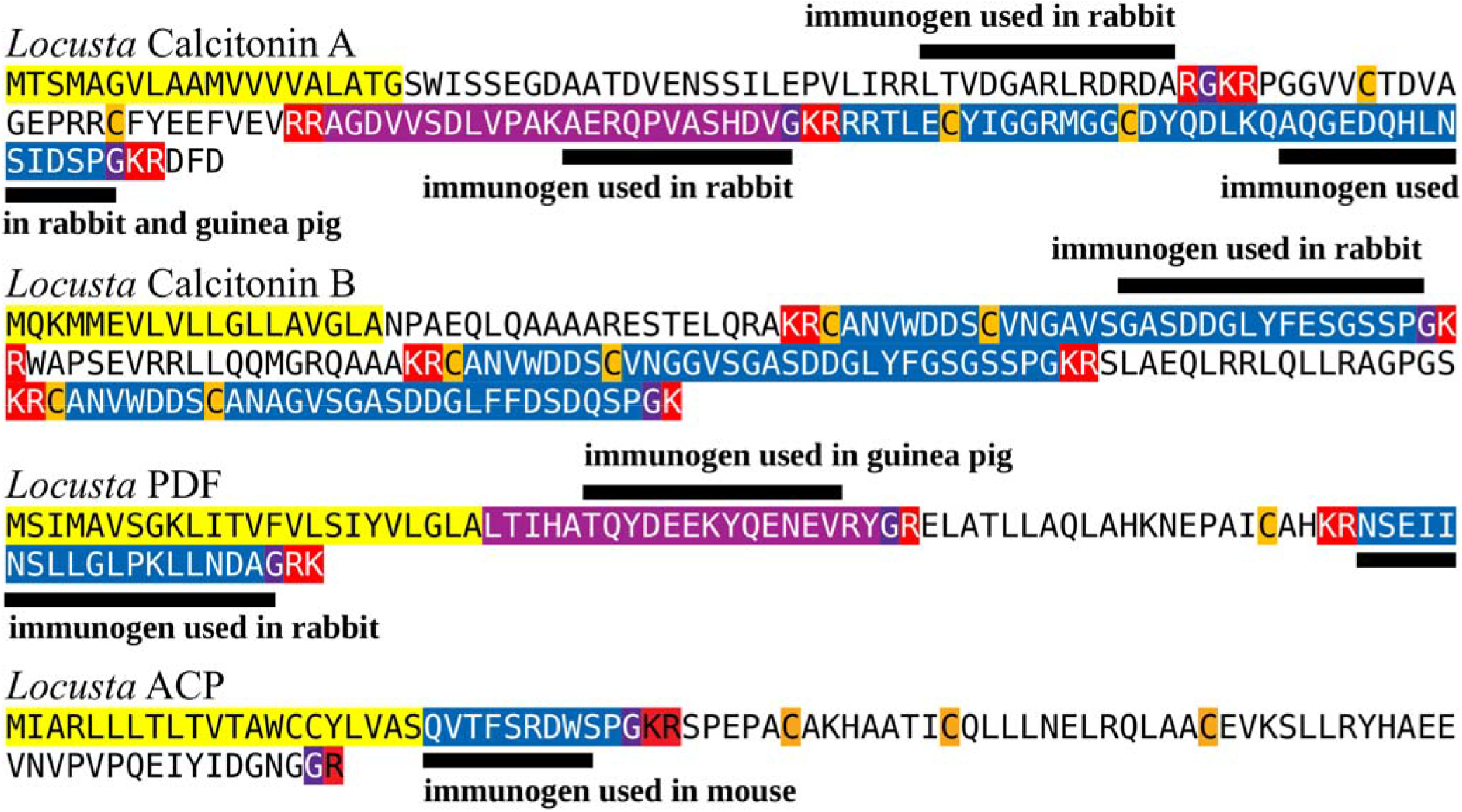
Precursors for the neuropeptides calcitonin A, calcitonin B, PDF and ACP from *Locusta migratoria*. Predicted signal peptides are indicated in yellow, known neuropeptides in blue, proteolytic cleavage sites and amino acid residues subsequently removed by carboxypeptidase during processing are indicated in red, while glycine residues that are expected to be transformed into C-terminal amides are in deep purple. Two other C-terminally amidated neuropeptides that are predicted from the calcitonin A and PDF precursors are highlighted in light purple. Several rabbits, two guinea pigs and a mouse were used to generate specific antisera to epitopes that are indicated by bold under- or overlines.

QAQGEDQHLNSIDSPamide, LTVDGARLRDRDA and GASDDGLYFESGSSPamide were conjugated to bovine serum albumin (BSA) with difluorodinitrobenzene as described by Tager (1976) and dialyzed against phosphate buffered saline pH 7.4. The other conjugates were prepared in a two step reaction. First BSA was allowed to react with sulfo-SMCC (Sulfosuccinimidyl-4-[N-maleimidomethyl]cyclohexane-1-carboxylate). After separating the derivatized BSA from unreacted reagent using gel filtration on Sepharose P50 it was then allowed to react with the peptides through the added cysteine residues. Those conjugates were separated from unreacted excess peptide by gel fitration on Sepharose P50. These conjguates were sent to Pineda Antikörper-Service (Berlin, Germany) for antiserum production.

The primary rabbit antiserum to PDF was diluted 1: 4,000, the PDF monoclonal antibody 1:10, the rabbit antisera to calcitonin A and B 1:10,000, the guinea pig antiserum to calcitonin A 1:2,000, the guinea pig antiserum to the PDF precursor 1:10,000 and the mouse ACP antiserum 1:1,000 to 1,2000. Other antisera were diluted as previously described. Secondary antisera were Alexa488-labeled goat anti-rabbit IgG, Cy3-labeled goat anti-guinea pig IgG and Cy3-labeled goat anti-mouse IgG (Jackson Immunoresearch Europe Ltd, Ely, UK) and were diluted 1:1,000 to 1:2,000.

The immunohistological and *in situ* hybridization procedures as well as the method to simultaneously perform both procedures on the same tissue was recently described in detail elsewhere (Veenstra, 2021).

### Affinity columns and antiserum purification

PRAESTO® NHS90 Agarose resin was purchased from Purolite Ltd (Liantrisant, Wales, UK, CF72 8LF) and affinity columns containing about 1 ml of resin were produced as follows. After washing out the isopropanol with 5 ml of icecold 1 mM HCl the resion was equilibrated in the coupling buffer that consisted of 0.5 M NaCl and 0.2 M NaHCO3 pH 8.3, after which the column was loaded with 1.2 ml of coupling buffer containing about 1.2 mg of peptide and the flow stopped for 30 min. Unreacted NHS was inactivated by incubating the the column with 1 M TrisHCl pH 8.5 for 30 minutes. Next followed three cycles of washing the column with 8 ml of 100 mM NaCH3CO2 and 500 mM NaCl (wash buffer) followed by 8ml coupling buffer. The column was then equilibrated with 10 ml PBS containing 0.1 % NaN3. Four different columns were made this way with *Gryllus* PDF, CAERQQVASHDVamide, GASDDGLYFESGSSPamide and CAQGEDQHLNSIDSPamide.

These columns were not used to affinity purify the respective antisera, but rather to remove cross-reacting IgGs. One ml of the rabbit PDF antiserum diluted 1:100 with PBS containing 0.1% NaN3 was passed through the HDVamide column. The other antisera were raised for this study and as large quantities were available 1 ml of antisera diluted 1:1 with PBS was used. HDVamide antiserum was passed through the PDF affinity column, rabbit calcitonin A antiserum through the calcitonin B affinity column and the rabbit calcitonin B antiserum through the calcitonin A affinity column. Eluted preabsorbed antisera were collected and tested at the same time and in the same final concentrations as raw antisera.

### RT-PCR

Tissues were dissected under saline and either frozen at -75 °C or immediately processed. Total RNA was extracted using mini spin columns from Macherey-Nagel GmbH & Co. KG, Dueren, Germany. One μg of RNA was reverse transcribed in a 20 μl reaction using Moloney Murine Leukemia Virus Reverse Transcriptase (New England Biolabs, Evry, France) and random primers and cDNA obtained amplified using Q5® DNA Polymerase (New England Biolabs) with gene specific oligonucleotide primers (Table 1).

**Table 1.**
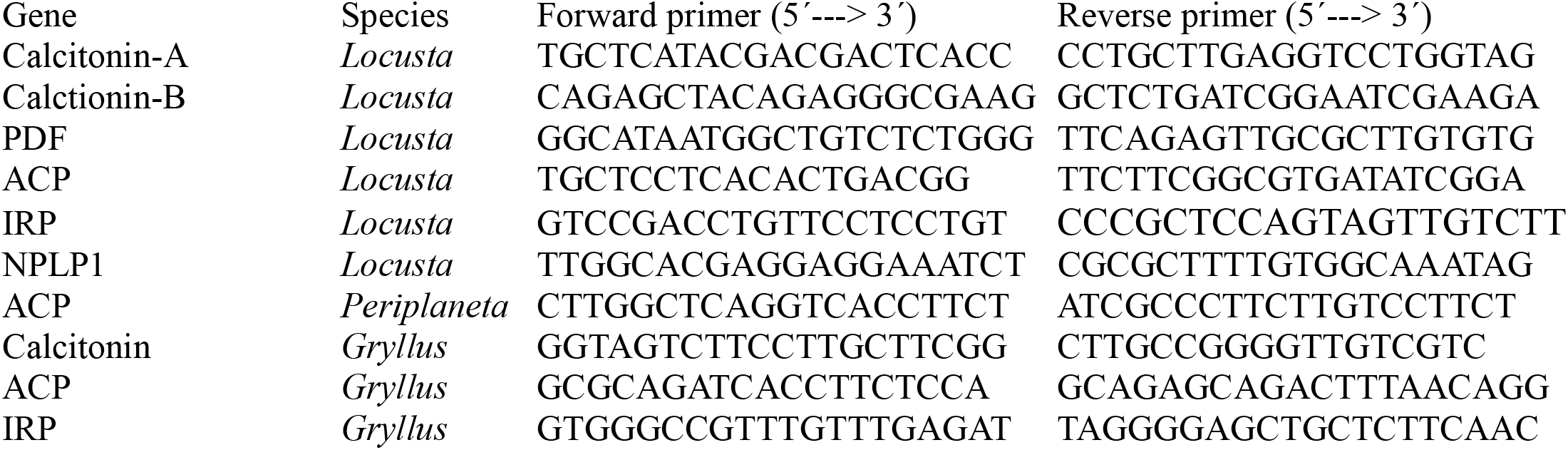
Oligonucleotide primers used for RT-PCR.

### In situ hybridization probes

cDNA’s coding partial calcitonin A, calcitonin B, PDF and NPLP1 precursors were amplified from cDNA from abdominal ganglia, midguts and brains respectively using the same gene specific primers (Table 1). The PCR products were gel purified and used to prepare digoxigenin labeled anti-sense DNA probes using PCR with digoxigenin-dUTP in a one sided PCR reaction using the reverse primers.

### Bioinformatics

Construction of coding sequences for neuropeptide precursors were done as described in detail elsewhere (Veenstra, 2020) using publicly available short read archives (SRAs) listed in the supplementary data. In order to find *Locusta* nervous system specific transcripts that might be recognized by the calcitonin antisera two CNS transcriptomes (Wang et al., 2015; Zhang et al., 2017) were downloaded from NCBI (https://www.ncbi.nlm.nih.gov/Traces/wgs/?term=tsa). The python2 script DNA2protein (https://zenodo.org/record/34023#.YIkRayaxVN0) was used to obtain the largest protein sequences coded by these transcripts and these were searched for the presence of sequences containing a PGKR subsequence. Those sequences were then inspected as to the likelihood they might represent a neuropeptide precursor.

## Results

### Antiserum specificity problems and solutions

Initial experiments using the guinea pig calcitonin A antiserum revealed neurons and neuroendocrine cells in the *Locusta* central nervous sytem that had previously been shown to react with a rabbit antiserum to PDF. These are four weakly immunoreactive neurons in the lower protocerebrum that project to the tritocerebrum (Homberg et al., 1991) and three bilateral pairs of ventral neuroendocrine cells in abdominal ganglia that have been called AP1 and are located posteriorly and medially in the abdominal neuromeres 5, 6 and 7 (Persson et al., 2001). *In situ* hybridization confirmed expression of the calcitonin A gene in the AP1 neuroendocrine cells (Fig. 2a) but failed to reliably detect expression in the four protocerebral neurons; on just one occasion an extremely faint signal was observed, but it could not be reproduced. Absence of a consistent *in situ* hybridization signal in the latter might reflect a much lower expression of the calcitonin A gene in the neurons then in neuroendocrine cells.

**Figure 2.**
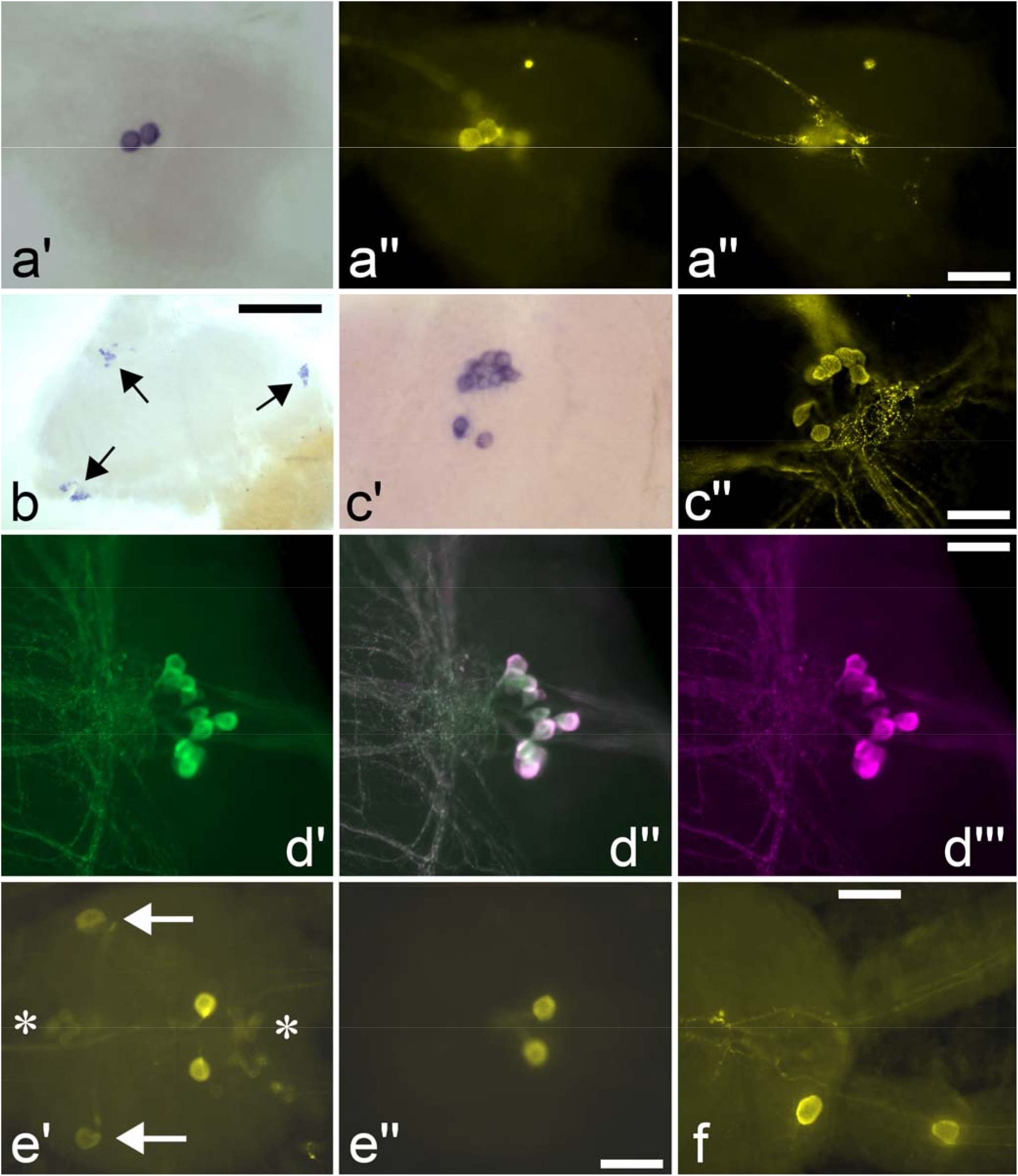
Selected illustration of immunoreactivity and *in situ* hybridizations localizing cells expressing PDF and calcitonin A in *Locusta migratoria*. Calcitonin A *in situ* hybridization signal (a’) in the AP1 cells is co-localized with calcitonin A immunoreactivity (a’’), the out of focus immunoreactive signal is due to axonal arborization of these cells (a’’’). Three groups in the optic lobe (arrows) are labeled by anti-sense PDF DNA (b), the same cells are also immunoreactive with PDF antiserum as illustrated for the cell group at the anterior edge of the medulla (c’, c’’). Rabbit antiserum to PDF (d’) labels the same neurons and axons (d’’) as guinea pig antiserum to the PDF precursor (d’’’). Rabbit antiserum to HDVamide labels not only the calcitonin A expressing AP1 neuroendocrine cells (e’) but also the AP2 neurons (axons) as well as several smaller cells (asterisks), however after passage of this antiserum through a PDF affinity column only the AP1 cells are labeled (e’’). Occasionally some AP1 neuronendocrine cells are located outside the ganglion proper and are found inside the connectives (f). Scale bars 100 μm (a, c, d & e) and 400 μm (b).

It seemed of interest to have an antiserum to different parts of the calcitonin A precursor to confirm that the immunoreactive protocerebral neurons do indeed express the calcitonin A gene. A rabbit antiserum to HDVamide, a second potential neuropeptide that might be produced from the calcitonin A precursor (Fig. 1), labeled all the calcitonin A immunoreactive neurons described above. Interestingly, this antiserum also strongly labeled the PDF neurons in the optic lobes and much more weakly the AP2 PDF-immunoreactive neuroendocrine cells of the abdominal ganglia (Persson et al., 2001) as well as what might be periviscerokinin cells in the abdominal ganglia (Eckert et al., 1999). The AP2 neurons have been described as dorsal neurons located anteriorly and laterally in neuromeres 3 through 7 (Persson et al., 2001), while the optic lobe neurons are two groups located at the posterior dorsal and posterior ventral of the lamina and a third group at anterior edge of the medulla (Homberg et al., 1991). However, *in situ* hybrdization with an antisense PDF probe only labeled the three groups of PDF neurons in the optic lobes (Fig. 2b,c) but labeled neither the AP1 nor the AP2 cells.

The immunoreactivity of all antisera was suppressed by preincubation with the peptides used to produce them and hence their immunoreacitivity was considered specific. As the PDF antiserum is widely used and considered reliable for detecting PDF immunoreactivity and at least the abdominal AP1 neuroendocrine produce calcitonin A as confirmed by the *in situ* hybridization experiments, it looked like these cells co-express calcitonin A and PDF. If so, the other calcitonin A and PDF immunoreactive neurons might also co-express these same two neuropeptide genes, but perhaps in different ratios. It seemed plausible that the optic lobe neurons might produce too little of calcitonin A to be detectable by the guinea pig antiserum to calcitonin A, since this antiserum gave a relatively weak signal.

These considerations led to the production of two other antisera, one guinea pig antiserum to a different part of the *Locusta* PDF precursor and a rabbit antiserum to calcitonin A. The guinea pig PDF precursor antiserum labeled fine PDF axons from the optic lobe neurons as intensely as the rabbit PDF antiserum (Fig. 2d), but it never labeled the four protocerebral neurons, nor the abdominal AP1 or AP2 neurondocrine cells. The labeling of this guinea pig antiserum was completely identical to that of the mouse monoclonal antibody to *Drosophila* PDF, that neither labels the AP1 nor the AP2 neurons. This raised doubts as to the specificity of the rabbit PDF antiserum. The rabbit calcitonin A antiserum labeled not only the same cells as the guinea pig calcitonin A antiserum, but also several other neurons, suggesting it cross-reacted with other neuropeptides. The most prominent of these neurons are neuroendocrine cells in the pars intermedia and the pars lateralis of the brain (Fig. 3). Nevertheless, it did not label the PDF neurons in the optic lobe, therefore, it seemed unlikely that the latter would express calcitonin A.

**Figure 3.**
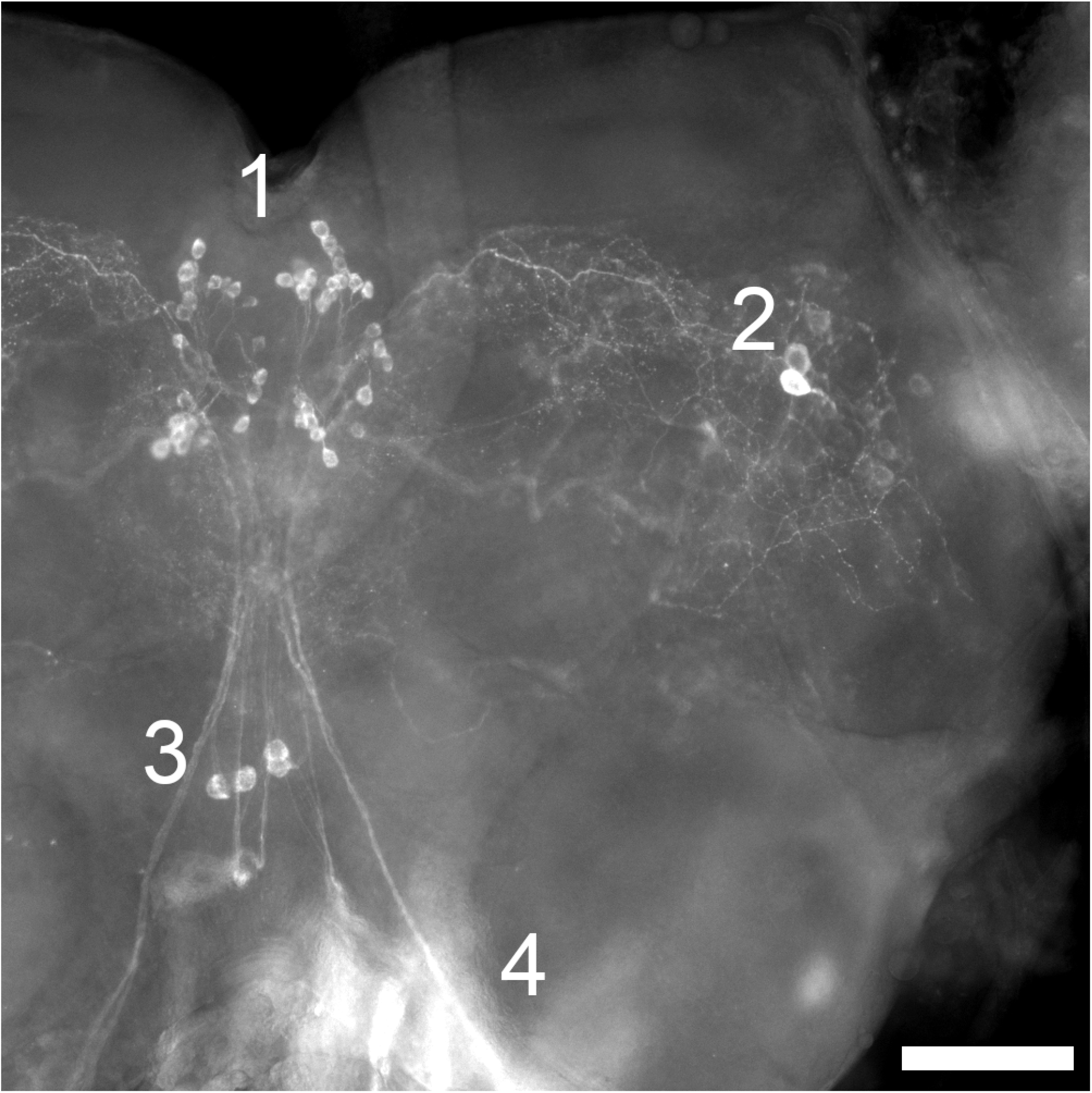
Calcitonin A immunoreactivity in the brain of *Locusta migratoria*. Three different cell types are visible in this picture, neuroendocrine cells in the pars intermedia (1) that project to the corpora cardiaca (a small part of which is out of focus in 4), lateral peptidergic interneurons (2) of which the largest two have extensive arborizations that are visible in the picture and the four neurons in the lower protocerebrum (3) that are the only ones that produce calcitonin A. Scale bar 200 µm.

The intron-exon boundaries of the *Locusta* calcitonin A gene are such that this gene could be alternatively spliced and produce mRNA species coding for three different precursors, the one illustrated in Fig. 1, or two others that lack either the calcitonin A or the HDVamide coding exon (Fig. S1). As other insect species have a single calcitonin gene from which mRNA’s coding precursors for calcitonin A an B are alternatively spliced (Veenstra, 2014), alternative splicing of the calcitonin A gene seemed a realistic possibility. Analysis of publicly available *Locusta* transcriptome SRAs did not find any evidence for alternatively spliced transcripts from this gene. However, as such transcripts might have escaped detection, yet another antiserum was produced, this one against RDRDA, a fragment of the precursor that would be present in all three hypothetical transcripts (Fig. S1). This antiserum strongly labeled the same cells as the guinea pig calcitonin A antiserum. Since despite its potency it did not label the PDF neurons in the optic lobe, this meant that these neurons do not express the calcitonin A gene and thus can neither produce HDVamide, raising also doubts as to the specificity of the HDVamide antiserum.

The similarity between the dipeptides DAmide and DVamide, the C-termini of PDF and HDVamide respectively, suggested that this might be the reason these antisera cross-react. After passage over a PDF affinity column, the HDVamide antiserum only recognized the same cells that are labeled by the guinea pig calcitonin A and rabbit RDRDA antisera and no longer labeled the PDF neurons in the optic lobe or the AP2 neurons (Fig. 2e). Interestingly, the immunoreactivity of this PDF preabsorbed HDVamide antiserum is noticeably stronger than the crude antiserum used at the same concentration. The PDF antiserum that was passed through a HDVamide affinity column no longer recognized the four protocerebral neurons nor the abdominal AP1 and AP2 neuroendocrine cells, but was still as immunoreactive toward the optic lobe PDF neurons. These results show that the PDF and HDVamide antisera cross-react and also how this cross-reactivity can be remediated. *Drosophila melanogaster* is a species that lacks genes coding calcitonin A or HDVamide. As expected the crude, but not the PDF preaborbed HDVamide antiserum labels its PDF neurons. These results suggested that the PDF immunoreactive material in the AP2 neurons might similarly be a C-terminal DXamide. Of all the knonwn *Locusta* neuropeptide genes NPLP1 looked like a plausible candidate to be expressed in the AP2’s. A previously missed coding exon of this gene encodes FLGVPPAAADYamide (Fig. S2), but *in situ* hybridization of an NPLP1 antisense probe did not label the AP2 cells (Fig. S3).

Both the protocerebral calcitonin A neurons and the abdominal AP1 neuroendocrine cells have been described in detail elsewhere so there is no need to do that here (Homberg et al., 1991; Persson et al., 2001). Nevertheless, two details are worth adding to the description of the AP1. On several occasions, these neurons were found outside the ganglia (Figs. 2f). This suggests that these neurons do not originate within the ganglion but migrate there during embryogenesis, as is also known for other arthropod and vertebrate neuroendocrine cells (de Velasco et al., 2007; Schwanzel-Fukuda and Pfaff, 1989). Interestingly, there seems to be a gradient in expression of the calcitonin A gene in these cells. When the alkaline phosphatase reaction is stopped relatively quickly, one often observes that the cells in neuromere 5 have the least signal and those in neuromere 7 the most (Fig. 4).

**Figure 4.**
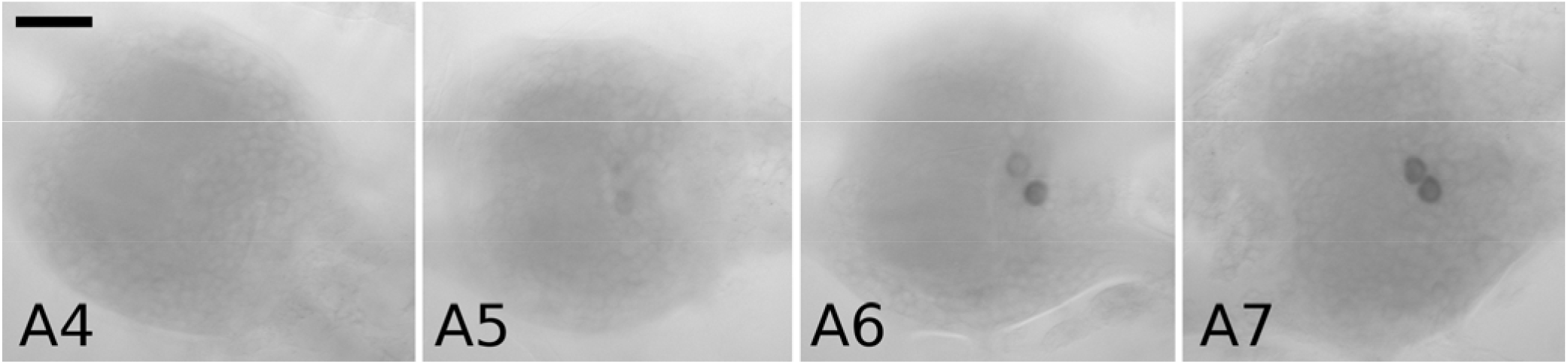
Calcitonin A *in situ* hybridization signal in abdominal ganglia of *Locusta* corresponding to the abdominal neuroemeres 4, 5, 6 and 7 (A4, A5, A6, A7). The alkaline phosphatase reaction was stopped very quickly to illustrate the difference in expression in these ganglia. In A5 signal is almost absent, in A6 it is very weak, and in A7 it is much stronger. Scale bar 100 μm.

Immunoreactivity with the different specific calcitoinin antisera in *Schistocerca gregaria* is very similar to that in *Locusta*. The major difference is that *Schistocerca* has an additional bilateral calcitonin immunoreactive neuroendocrine cell in abdominal neuromere 4. Minor differences are that HDVamide antiserum does not recognize homologs of the AP2 abdominal neurons and that the PDF preabsorbed HDVamide antisera barely recognizes the abdominal AP1 neurons, while the RDRDA antiserum is not immunoreactive in this species. The latter two observations are not surprising, as the the sequence of the *Schistocerca* Calcitonin A precursor differs significantly from *Locusta* in the peptide sequences used to make these antisera (Fig. S4).

None of the cricket genome assemblies, for *Laupala kohalensis, Gryllus bimaculatus, Teleogryllus occipitalis* and *T. oceanicus* (Blankers et al., 2018; Pascoal et al., 2019; Ylla et al., 2020; Kataoka et al., 2020), contain sequences coding for a calcitonin B ortholog or its putative receptor, while they do have genes encoding calcitonin A (Fig. S4) and its putative receptor. In *G. bimaculatus in situ* hybrdization labels cells that appear homologs of the *Locusta* AP1’s, with additional cells in the terminal abodminal ganglion (Fig. S6). Such cells were also found in the abdominal ganglia of the cockroach *Periplaneta americana* using the rabbit calcitonin A antiserum (Fig. S7), as in the locust these cells were some times found outside the ganglia.

The antiserum to calcitonin B labels enteroendocrine cells that are the same as those labeled by an antisense probe for the calcitonin B gene (Fig. S8). These cells are most prominent in the gastric caeca, particularly in the smaller posterior ones, but they are also present in midgut proper (Fig. 5a,b,c). However, calcitonin B is not the only peptide recognized by this antiserum. When tested on the central nervous system it reacts with several cell types including those that are recognized by the guinea pig calcitonin A and RDRDA antisera, but also what seemed to be the same neuroendocrine cells in the *pars mediana* and *pars lateralis* that are recognized by the rabbit calcitonin A antiserum. After passing the calcitonin B antiserum through a calcitonin A affinity column the antiserum no longer recognized any neurons or neuroendocrine cells the central nervous system, but this did not affect its immunoreactivity toward the enteroendocrine cells (Fig. 5d,e). Passing the rabbit calcitonin A antiserum through a calcitonin B affinity column similarly removed the immunoreactivity toward the neuroendocrine cells of *pars intermedia* and the *pars lateralis*. This suggested that the two antisera recognized the same peptide(s).

**Figure 5.**
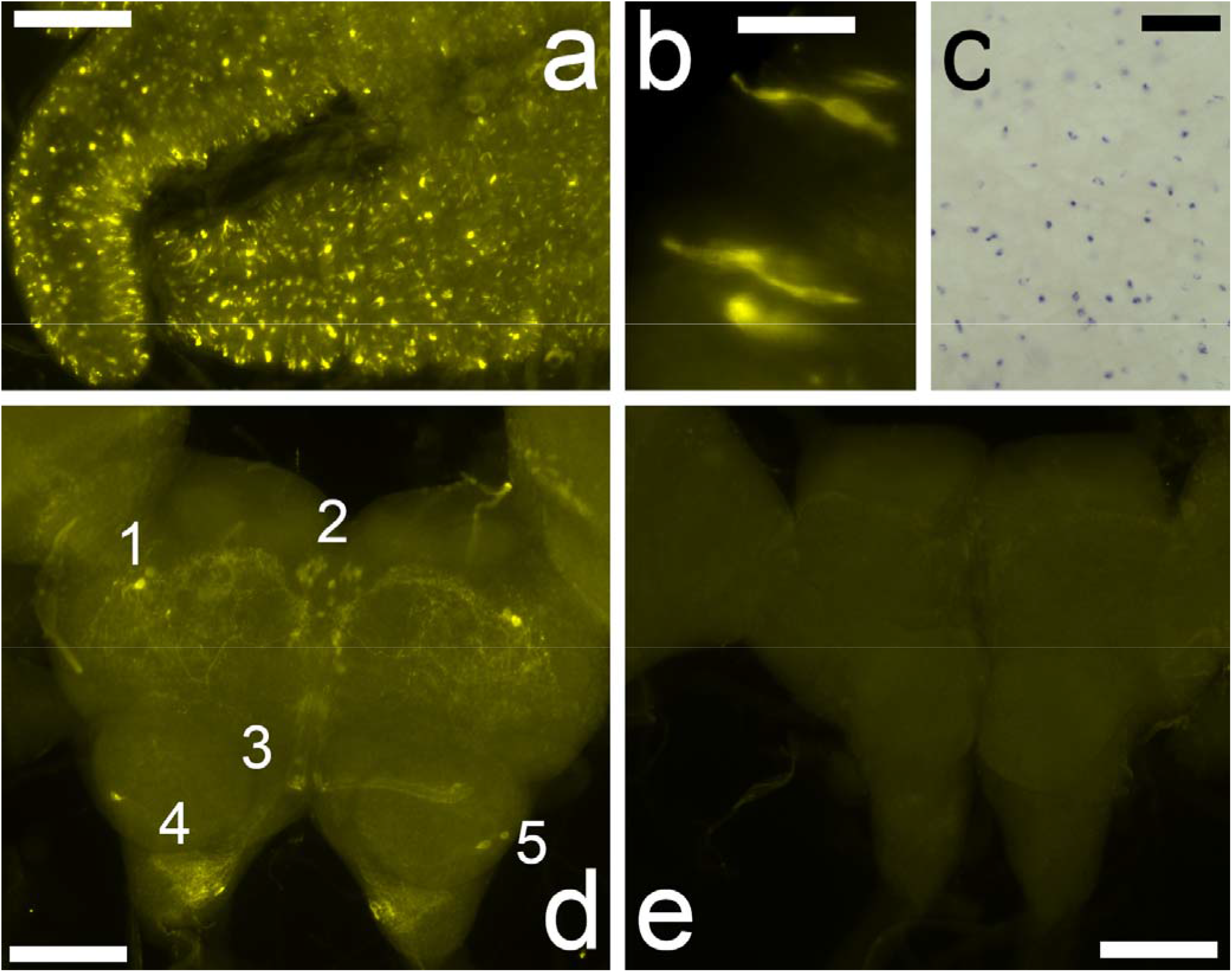
Calcitonin B immunoreactivity in the intestin is specific, but in the brain is due to cross-reactivity. Calcitonin B immunoreactivity is particulary abundant in the posterior caeca (a) and present in cells that have the typical morphology of enteroendocrine cells (b). *In situ* hybridization with an antisense calcitonin B probe labels numerous cells in the caeca as well as in the midgut proper (c). Calcitonin B immunoreactivity in the brain (d) is unspecific, since it is completely abolished after the antiserum is passed through a calcitonin A affinity column (e). 1: neuroendocrine cells of the *pars lateralis*; 2: peptidergic interneurons of the *pars intermedia*; 3: very weakly labeled and largely out of focus the four calcitonin calcitonin A neurons; 4: the arborizations of the calcitonin A neurons in the tritocerebrum; 5: unidentified immunoreactive neurons in the deutocerebrum. Scale bars 50 μm (b), 100 μm (c, d, e) or 400 μm (a).

Although *Locusta* calcitonins A and B are paralogs, the sequence similarity between the two peptides used to raise antisera is limited to the C-terminal dipeptide SPamide (Fig. S5), suggesting that this C-terminal dipeptide might be responsible for the cross-reactivity between these antisera. In order to identify the calcitonin immunoreactive peptide in the brain neuroendocrine cells I used two *Locusta* central nervous system transcriptomes (Wang et al., 2015; Zhang et al., 2017) to look for transcripts that can code for a PGKR sequence. Such transcripts might correspond to precursors coding neuropeptides with a C-terminal Proline-amide. Of the 52 and 103 sequences found, two coded neuropeptide precursors, *i*.*e*. calcitonin A and ACP. Like the *Locusta* calcitonins *Locusta* ACP also has a C-terminal SPamide sequence. Double labeling with the mouse ACP and rabbit calcitonin antisera showed that all three antisera label the same neuroendocrine cells, both in the *pars mediana* and the *pars lateralis* of the brain. Signal obtained from *in situ* hybridization with an antisense ACP probe is present in the same cells that are immunoreactive with the ACP antiserum (Fig. 6), thus confirming the specificity of the ACP antiserum. These cells were very recently also described using a rabbit antiserum to ACP (Hou et al., 2021). The data obtained for the expression of these four *Locusta* neuropeptide genes, calcitonins A and B, ACP and PDF, are summarized in Table 2.

**Table 2.** Results of the various antisera and *in situ* hybridization experiments. AP1 and AP2, the PDF-immunoreactive AP1 neurons described by Persson et al., 2001; Girardie, the four protocerebral neurons initially described by Girardie (1970); LNCS, lateral neurosecretory cells, those that are located in the *pars lateralis*; MNSC, median neurosecretory cells, those that are located in the *pars intermedia*; Optic lobe refers to three PDF immunoreactive neurons in the optic lobe. +++ strong signal; (+)? dubious signal; - no signal; n.t., not tested. Preabsorbed in this context means after passage through an affinity column of said peptide.

|  | Cell type with neuropeptide it expresses |  |  |  |  |  |  |  |
| --- | --- | --- | --- | --- | --- | --- | --- | --- |
|  | Optic lobe<br>PDF | LNSC<br>ACP | MNSC<br>ACP | Girardie<br>Calcitonin A | AP1<br>Calcitonin A | AP2<br>unknown | other<br>unknown | Enterocyt<br>Calcitonin |
| rb PDF | +++ | - | - | ++ | +++ | ++ | - |  |
| rb PDF HDVamide-column | +++ | - | - | - | - | - | - |  |
| monoclonal PDF | +++ | - | - | - | - | - | - |  |
| gp PDF-precursor | +++ | - | - | - | - | - | - |  |
| PDF *in situ* hybridisation | +++ | - | - | - | - | - | - |  |
| gp Calcitonin A | - | - | - | ++ | +++ | - | - |  |
| rb Calcitonin A | - | - | - | ++ | +++ | - | - |  |
| rb HDVamide | +++ | - | - | ++ | +++ | ++ | - |  |
| rb HDVamide PDF-column | - | - | - | ++ | +++ | - | - |  |
| rb RDRDA antiserum | - | - | - | ++ | +++ | - | - |  |
| rb Calcitonin A | - | +++ | +++ | ++ | +++ | - | + |  |
| rb Calcitonin A Calc.B-column | - |  |  | ++ | +++ | - | - |  |
| rb Calcitonin B | - | +++ | +++ | ++ | +++ | - | + |  |
| rb Calcitonin B Calc.A-column | - | - | - | - | - | - | - |  |
| Calcitonin A *in situ* | - | - | - | (+)? | +++ | - | - |  |
| Calcitonin B *in situ* | - | - | - | - | - | - | - |  |
| mouse ACP | - | +++ | +++ | - | - | - | - |  |
| ACP *in situ* | - | +++ | +++ | - | - | - | - |  |

**Figure 6.**
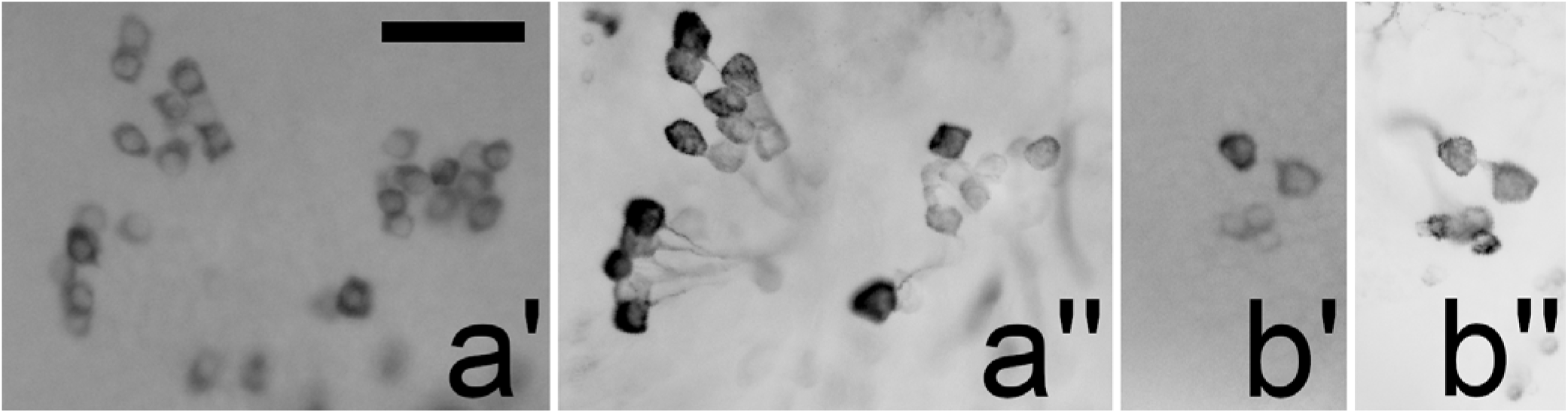
ACP-immunoreactive cells in the brain (a’’,b’’) are the same as those labeled by *in situ* hybridization with an ACP antisense probe (a’,b’), both the neuroendocorine cells in the *pars intermedia* that project to the corpora cardiaca (a) and the interneurons in the *pars lateralis* (b). Scale bar 100 μm.

Three major neuroendocrine cell types in the *pars intermedia* of *Locusta* that, like the ACP cells, project to the corpus cardicaum, are known to produce neuroparsin, IRP or both a CRF-like diuretic hormone and ovary maturating parsin (Bourême et al., 1989; Goltzené et al., 1992; Patel et al., 1994; Tamarelle et al., 2000). Combined *in situ* hybridization with anti-sense IRP probe showed that the ACP cells are different from the IRP producing cells (Fig. S9), while double labeling with rabbit antisera to *Locusta* CRF-like diuretic hormone and neuroparsin showed that ACP is neither co-localized with those peptides. The *pars lateralis* has peptidergic neurons that produce corazonin, allatostatins A and B, leucokinin and ion-transport peptide. Rabbit antisera to these peptides reacted with neurons that are different from the ACP immunoreactive neurons.

In other insect species ACP had so far only been found in interneurons and never in neuroendocrine cells of the *pars intermedia* projecting to the corpora cardiaca (Hansen et al., 2010; Patel et al., 2014). Preliminary experiments showed that *Schistocerca gregaria* and *Gryllus bimaculatus* also have neuroendocrine cells in the *pars intermedia* that project to the corpora cardiaca, but in the cockroach *Periplaneta americana* ACP immunoreactivity and in situ hybridization signal is only found in the *pars lateralis* of the brain.

## Discussion

This study describes the expression pattern of *Locusta* calcitonins and corrects that of PDF and illustrates the danger of relying on a single antiserum to determine the expression of a neuropeptide. It also shows that the abdominal neuroendocrine cells producing calcitonin A seem to be conserved in at least some Polyneoptera, while in Orthoptera the expression of ACP is remarkably different between from that in other insect species.

Specificity of antisera to neuropeptides is often difficult to ascertain. The results reported here show that four previously identified PDF immunoreactive neurons in the lower protocerebrum and six PDF immunoreactive neuroendocrine cells in the abdominal ganglia in *Locusta migratoria* do not produce PDF but calcitonin A, while a second type of PDF-immunoreactive neuroendocrine cells in the abdominal ganglia neither produce PDF. Although the sequence similarity between HDVamide and PDF is limited to the extreme C-termini of these peptides, HDVamide and NDAamide, this appears to be sufficient for crossreactivity. In hindsight it is probably not too surprising, since FMRFamide antisera also only need the RFamide or RYamide C-termini to recognize neuropeptide producing neurons in immunohistology (*e*.*g*. Veenstra and Schooneveld, 1984; Veenstra, 1984; Veenstra and Khammassi, 2017). Indeed, in retrospect the FMRFamide and vasopressin immunoreactivity of neuroendocrine cells in the suboesophageal ganglion of *Locusta* (Veenstra, 1984) must be due to the presence of tryptopyrokinins, with their C-terminal R(I/V)amides sequences, suggesting that for the FMRFamide antisera RXamide may be sufficient for immunoreactivity. It is well known that electrostatic interactions are much stronger than hydrophobic interactions. The examples of RFamide and HDVamide antisera show that the positive or negative charge of a single amino acid residue can be sufficient for antigen binding in immunohistology.

Whereas the cross-reactivity of the PDF and HDVamide antisera can be rationalized by assuming a major contribution of charged amino acid residues to antigen-antibody interactions, such charged amino acid residues are absent from the SPamide C-termini of the calcitonins A and B. Perhaps it is the rigid conformation of the proline that is responsible for the relatively strong immunoreactivity of the calcitonin antisera toward this part of these peptides.

Although polyclonal antisera are often purified on affinity columns to remove non-specific antibodies from the serum, the antibodies with the highest affinity for the antigen are often not recovered from such columns. Here I used affinity columns, not to recover the specific antibodies, but to remove the cross-reacting ones, thus likely recovering the high affinity antibodies. After passage through the PDF affinity column, the HDVamide antiserum yielded better immunoreactivity. This may be explained by assuming that the antiserum contains large numbers of lower affinity cross-reacting antibodies that compete with the less abundant but more specific and higher affinity antibodies for binding to the antigen in the preparation. After the first wash the antibodies that are not fixed to the tissue are removed and during the following washes many of those lower affinity antibodies detach from the antigen and are subsequently lost, leading to a weaker signal than when these less specific antibodies have been previously removed on the affinity column.

ACP was initially identified as an AKH-like peptide produced by the brain (Siegert, 1999). It was a surprise to see ACP being produced by neuroendocrine cells of the *pars intermedia* as in other insect species ACP seemed to be expressed in the lateral parts of the brain (Hansen et al., 2010; Patel et al., 2014). An extensive characterization of the *Locusta* neuropeptide gene was recently published.

After previously having shown that ACP is quickly upregulated when solitary nymphs are crowded, and thus transition to the gregarious phase (Hou et al., 2017), Hou and colleagues showed that the actions of this peptide on flight muscle is essential for both preparing and maintaining long distance flight in this species (Hou et al., 2021). It is worth recalling the partial characterized allatotropin from *Locusta*, as its description fits ACP remarkably well. This putative *Locusta* allatotropin shares with ACP its localization in the storage part of the corpora cardiaca, its molecular weight between 500 and 1500 and is digestability by both trypsin and chymotrypsin (Gadot et al., 1987). It will be interesting to know whether this is pure coincidence or whether some of the effects of ACP are achieved indirectly through juvenile hormone.

This work was initiated based on the hypothesis that four protocerebral neurons suggested to produce a diuretic hormone might express the calcitonin A gene and that calcitonin A might thus be a diuretic hormone. The cells in the inferior protocerebrum that produce calcitonin A are expected to be lightly stained by paraldehyde fuchsin. In the absence of other paraldehyde fuchsin positive cells in this part of the brain, the staining characteristics and their location identify these cells as the ones that were described by Girardie (1970) as possibly producing a diuretic hormone. However, these cells have no neuroendocrine release sites and can thus not release a diuretic hormone. Girardie described that their staining intensity changed noticeably depending on the relative humidity in which the insects were kept. On three occasions I have tried to confirm this by using diluted calcitonin A and HDVamide antisera, but no change in staining intensity was observed either in those cells or the abdominal AP1 neuroendocrine cells. Although, it still remains possible that the calcitonin A released from the abdominal neuroendocrine cells or calcitonin B released from the enteroendocrine cells may stimulate fluid secretion by the Malpighian tubules, these experiments do not support the hypothesis that the four protocerebral calcitonin A neurons are involved in the regulation of diuresis.

This raises the question of how electrocoagulation of those four neurons may have led to an increase in body weight as well as body length. A similar, and likely the same phenotype, was also observed on electrocoagulation of the median neuroendocrine cells of the *pars intermedia* (Cazal and Girardie, 1968), while an increase of body weight and body length has been observed in *Periplaneta* cockroaches under different as well as similar conditions. Thus, injection of the neuropeptide sNPF into female nymphs led to a significant increase in food consumption and weight within 24 hours (Zeng et al., 2021). This suggests that intensified feeding may be the proximate cause of the increase in body weight rather than the lack of a diuretic hormone. Such a significant increase in both food intake and weight gain was also found in adult males of the same species in which either the median neuroendocrine cells had been removed or in which the protocerebrum had been bisected. However, in those experiments it only occurred when these animals were kept under conditions of constant darkness, but not when the animals were exposed to a normal dark-light cycle. Importantly, when kept under constant darkness such animals also become arhythmic and hyperactive (Matsui et al., 2009). Thus in that experiment the animals that increase their body weight and length was associated with arhythmicity and hyperactivity.

Elegant experiments by Page in a different cockroach species have shown that the optic lobes entrain the circadian clock (Page, 1982). He disconnected the optic lobes from the midbrain and exchanged one of the optic lobes for one from an animal with a distinctly different free running cycle. Once the optic lobes in such animals reconnect with the midbrain each optic lobe independently imposes its rhythm on the circadian clock and such animals may develop activity patters consisting of two independent periodicities (Page, 1983), illustrating that the midbrain contains the clock that is entrained by the optic lobes. If bisectioning the brain completely abolishes the clock it is likely constituted by bilateral neurons that depend on mutual interactions. It is conceivable that extirpation of the median neurosecretory cells of the brain as well as cauterization of the calcitonin A neurons would also cut the connection between these clock neurons and inactivate it. Whereas cockroaches are active during darkness, locusts are active during the day. So perhaps destroying the circadian clock in *Locusta* by cauterizing either the median neurosecretory cells or the four protocerebral neurons similarly induced arhythmicity and hyperphagy.

## Acknowledgements

I am especially grateful to Julio Pineda for making the antisera; unfortunately his small company did not survive the covid19 pandemic. I also thank Justin Blau, Heinrich Dircksen, Geoffrey Coast and Josiane Girardie for graciously providing antibodies.

