## SupplementaryFigures for "Identification of cells expressing Calcitonins A and B, PDF and ACP in *Locusta migratoria* using cross-reacting antisera and *in situ* hybridization"

by Jan A. Veenstra

Contents:

List of SRAs used for analysis 2
Figure S1. Possible alternative splicing of *Locusta* calcitonin A 3
Figure S2. Improved prediction for the *Locusta* NPLP1 precursor 4
Figure S3. *In situ* hybridization of NPLP1 in *Locusta* abdominal ganglion 1 4
Figure S4. Sequence comparison of calcitonin A precursors 5
Figure S5. Sequence comparison of calcitonin A and B immunogens 5
Figure S6. Calcitonin A *in situ* hybridization in *Gryllus bimaculatus* 6
Figure S7. Calcitonin A immunoreactivity in *Periplaneta* 7
Figure S8. Calcitonin B and *in situ* hybridization in *Locusta* midgut 8
Figure S9. Combined insulin *in situ* and ACP immunohistology in *Locusta* brain 9
Figure S10. Sequence comparison of ACP precursors 10

**List of SRAs used for analysis:**

SRR167712, SRR3318257, SRR3318261, SRR3318262, SRR3318263, SRR3318264, SRR3318265, SRR3318266, SRR3318268, SRR3318269, SRR3318271, SRR3318305, SRR3318306, SRR4470191, SRR4470192, SRR4470193, SRR4470194, SRR4470195, SRR4470196, SRR4470197, SRR4470198, SRR4470199, SRR4470200, SRR4470201, SRR4470202, SRR5580661, SRR5580662, SRR5580663, SRR5580664, SRR5580665, SRR5580666, SRR5580667, SRR5580668, SRR5580669, SRR5580670, SRR5580671, SRR5580672, SRR5759362, SRR5759363, SRR5759364, SRR5759365, SRR5759366, SRR5759367, SRR5966534, SRR5966981, SRR5966984, SRR5967009, SRR5967010, SRR5967011, SRR6109303, SRR6109304, SRR6109305, SRR6109306, SRR6109307, SRR6109308, SRR6109309, SRR6109310, SRR6109311, SRR6109312, SRR6109313, SRR6109314, SRR6109315, SRR6109316, SRR6109317, SRR6109318, SRR6109319, SRR6109320, SRR6109321, SRR6109322, SRR6109323, SRR6109324, SRR6109325, SRR6109326, SRR8144091, SRR8144092, SRR8144093, SRR8144094, SRR8144095, SRR8144096, SRR8144097, SRR8144098, SRR8144099, SRR8144100, SRR8144101, SRR8144102, SRR8144103, SRR8144104, SRR8144105, SRR8494480, SRR8494481, SRR8494482, SRR8494483, SRR8494484, SRR8494485, SRR8643699, SRR8643700, SRR8643701, SRR8643702, SRR8643703 & SRR8643704.

RDRDA antiserum

*Locusta* Calcitonin A

MTSMAGVLAAMVVVVALATGSWISSEGDAATDVENSSILEPVLIRRLTVDGARLRDRDARGKRPGGVV
CTDVAGEPRRCFYEEFVEVRRAGDVVSDLVPAKAERQPVASHDVGKRRRTLECYIGGRMGGCDYQDLK
QAQGEDQHLNSIDSPGKRDFD

HDVamide antiserum

Calcitonin A antisera

Potential alternative transcripts:

RDRDA antiserum

HDVamide only

MTSMAGVLAAMVVVVALATGSWISSEGDAATDVENSSILEPVLIRRLTVDGARLRDRDARGKRPGGVV
CTDVAGEPRRCFYEEFVEVRRAGDVVSDLVPAKAERQPVASHDVG

HDVamide antiserum

RDRDA antiserum

Calcitonin A only

MTSMAGVLAAMVVVVALATGSWISSEGDAATDVENSSILEPVLIRRLTVDGARLRDRDARGKRRRTLE
CYIGGRMGGCDYQDLKQAQGEDQHLNSIDSPGKRDFD

Calcitonin A antisera

**Fig. S1.** The three different *Locusta* calcitonin A precursors that might be produced if the calcitonin A gene were alternatively spliced. The antisera to the three different peptides from this precursor would be able to distinguish between those three, if indeed this gene is alternatively spliced.

```
MRWCSLAALCAALAVAAMRPPQAGADPGDAVSVSVPGKRYVAALARNGALPLYGGWRKKQQRGDKRYTA
RYGKRADLDDEGGDDEPLTGVDALLEEIAATEQLRHLQLDALRRELDAQEAREEQAVLAAAAEAAAAEA
EDEEGPDAEADKRSVASLARAGALLPGKRNIAAMAKNGLLGPSGPVLLDGEGEKRSVGALARSGLLPQP
GRRAQDAGDDDLSLDSLMQQLYSEEEKRHIGTLARDYSLPSYGKRNLSLARSGGLSNVRYVTSKKDDSQ
PPADKRSLASLMRSRGPSPVEKRYLASLVRSHGLPYPLTKKEDDGPGEIKRNVGALARNWMLPSGKRAD
GDDQEVDKRYLASVLRQGRSDGFRQSSDSAQEAGHEEEKRHLGSLAKSGMAIHKKSSRSAGSDGQAFLQ
QQQQQQQEAGSKRAKRQAYLLPPAPPQSLAPGEFPMPVLQNTDDLSDYEDLLELMSGGLAAPEKRFLGV
PPAAADYGKRHIGALARLGWLPSFRAAPSRAGRSAGSRSGKRAYRSYPADGPWPSELQQA-
```

**Fig. S2.** Predicted sequence of the NPLP1 precursor. It contains an additional exon that encodes FLGVPPAAADYamide that might be recognized by the PDF and HDVamide antisera.

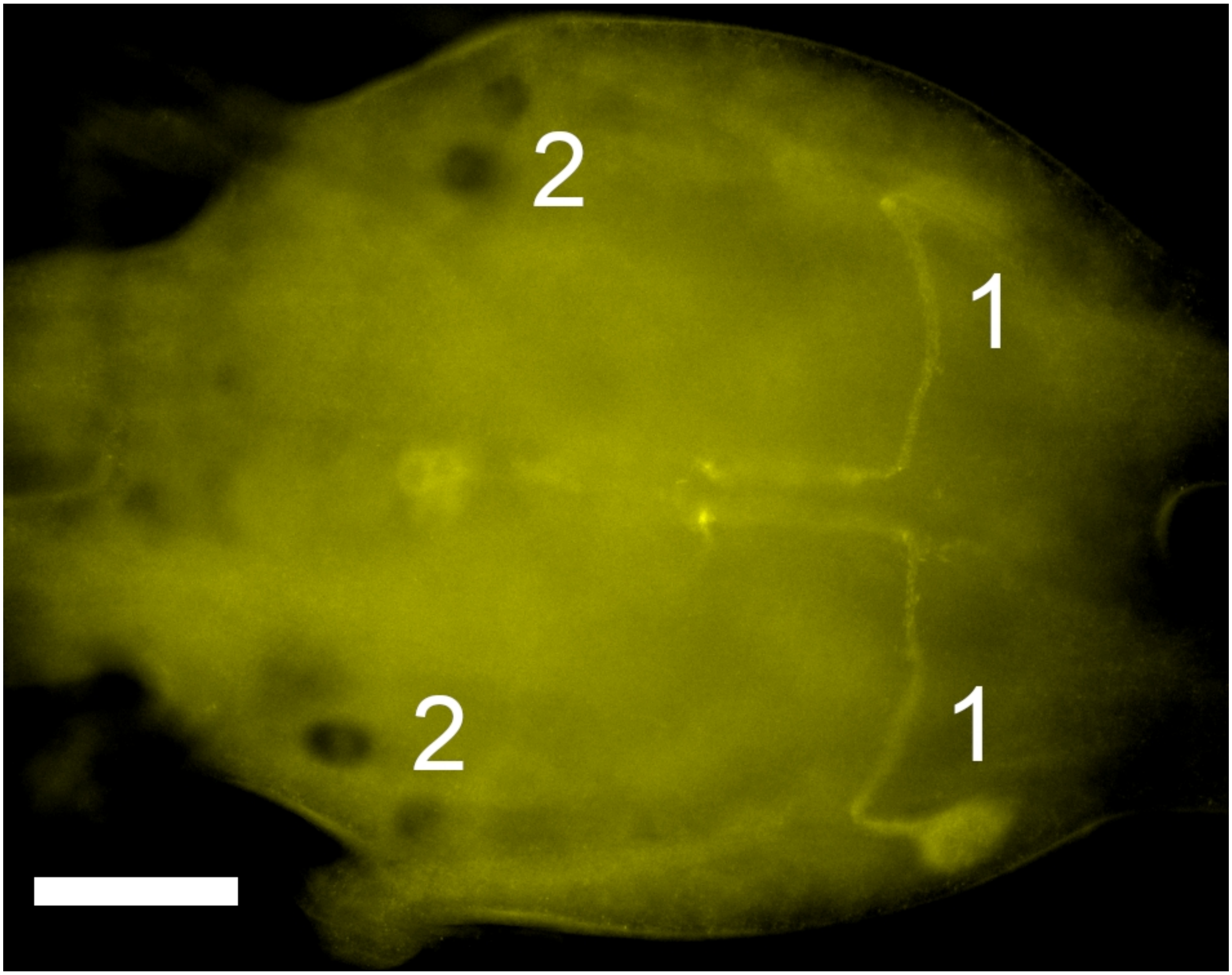


**Fig. S3.** First abdominal ganglion of *Locusta migratoria* showing the AP2 cells labeled by the HDVamide antiserum (1, the top AP2 cell is out of focus, as are the AP1 cells) and *in situ* hybrdization signal for the NPLP1 gene (black cells in two). Note that the AP2 cells do not appear to produce NPLP1.

# A

```
Locusta         MTSMAGVLAVMVAVVALATGSWISSEGEAATDV--ENSSILEPVLIRRLTVDGARLRDRD  58
Schistocerca    MTSTVGVLTAMVAAIALATGSWVPPDGDSARDV--ENSSILEPVFIRRLTIDGLRHRDYG  56
Gryllus            MASFTLVVFLA--SVAATMAMPRDISQSLIDSHMKHFEENRQALKLLKAIIDEMNSN  26
Periplaneta     MEWKREMTLVLYLLVVMATAAWASSKEIAQELMDSHIKSIQENRRTVRLLKNLLEDMDLN  31

Locusta         ARGKRRGGVVCTDVVGEPRRCFYEEFVEVRRAGDVVSDLVPAKAERQQVASHDVGKRRRT 118
Schistocerca    ----TRGGVVCTDVAGEPRRCFYEELVEMRRPEDVLNDLLSAKRDRQPLESHDVGKRRRT 114
Gryllus         ----------------------------------------------------I-EFVQKR  62
Periplaneta     ----------------------------------------------------M-ETVQKR  67

Locusta         LECYIGGRMG-GCDYQDLKQAQGEDQHLNSIDSPGKRDFD*                    157
Schistocerca    LECYIGGRMG-GCDYQDIKQAQGEDQHLNSIDSPGKRDLD*                    153
Gryllus         SVCYIGGGMGHNCEYGEALDSAFAARHLLGDDNPGKRSRLPLAPAGGPF*           111
Periplaneta     TSCLINAGLSHSCDNRDFIGAVEENKYWRSLDSPGKRRRRSSDNTQ*              113
```

# B

```
Locusta         AERQQVASHDV-amide
Schistocerca    RDRQPLESHDV-amide
```

# C

```
Locusta         LTVDGARLRDRDA
Schistocerca    LTIDGLRHRDYGTRG
```

**Fig. S4.** Sequence comparison of calcitonin A precursors from *Locusta migratoria, Schistocerce gregaria, Gryllus bimaculatus* and *Periplaneta americana.* **A:** The complete precursors. Note that the predicted mature locust calcitonin A sequences are virtually identical, but that they differ significantly from the other species. Yellow highlighting indicates the signal peptide, red amino acid residues are those that serve as substrate for convertase and carboxypeptidase, purple residues are glycine that are predicted to be transformed in C-terminal amides and orange ones cysteines that are predicted to form disulfide bridges. Blue residues correspond to the mature calcitonin A sequence, dark blue are those residues that are identical to the *Locusta* sequence. **B:** Comparison of the HDVamide sequences from *Locusta* and *Schistocerca*. Note that although the C-terminal is identical that the remainder of the peptide is significantly different, which probably explains that once the antibodies that specifically recognize the C-terminal (by passage through the PDF affinity column) antiserum to the *Locusta* peptide hardly recognizes the orthologous *Schistocerca* sequence. **C:** Sequence comparison of the third peptide used to make *Locusta* calcitonin A specific antiserum with the orthologous sequence from *Schistocerca*. Note that apart from the amino acids that are different, in *Locusta* the last amino acid residue, an alanine, has a C-terminal acid group confering an additional negative charge to this peptide, while in *Schistocerca* in stead there is an arginine residue conferring a positive charge to this peptide.

```
Calcitonin-A       AQGEDQHLNSIDSP-amide
Calcitonin-B      GASDDGLYFESGSSP-amide
```

**Figure S5.** Comparison of the C-terminal parts of *Locusta* calcitonins A and B used to make antisera. Note that the similarity between these two peptides is limited to the C-terminal dipeptide SP-amide.

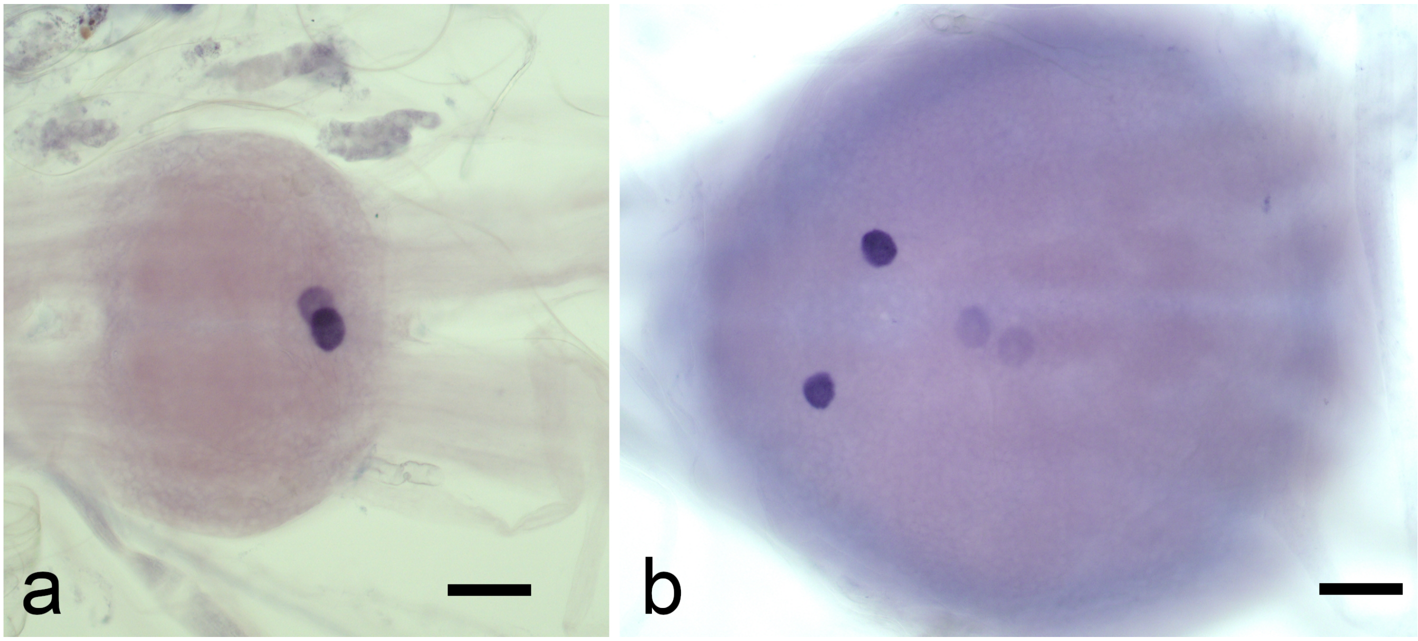


**Fig. S6.** Calcitonin A *in situ* hybridization in *Gryllus bimaculatus* abdominal ganglion 4 (a) and the terminal abdominal (b). As in *Locusta* there are two relatively large cells in the abdominal ganglia 2, 3 and 4. However, unlike *Locusta* there are also two large cells that show a strong signal in the terminal abdominal ganglia and two additional cells can be revealed when the reaction is let run longer. Scale bars 100 µm.

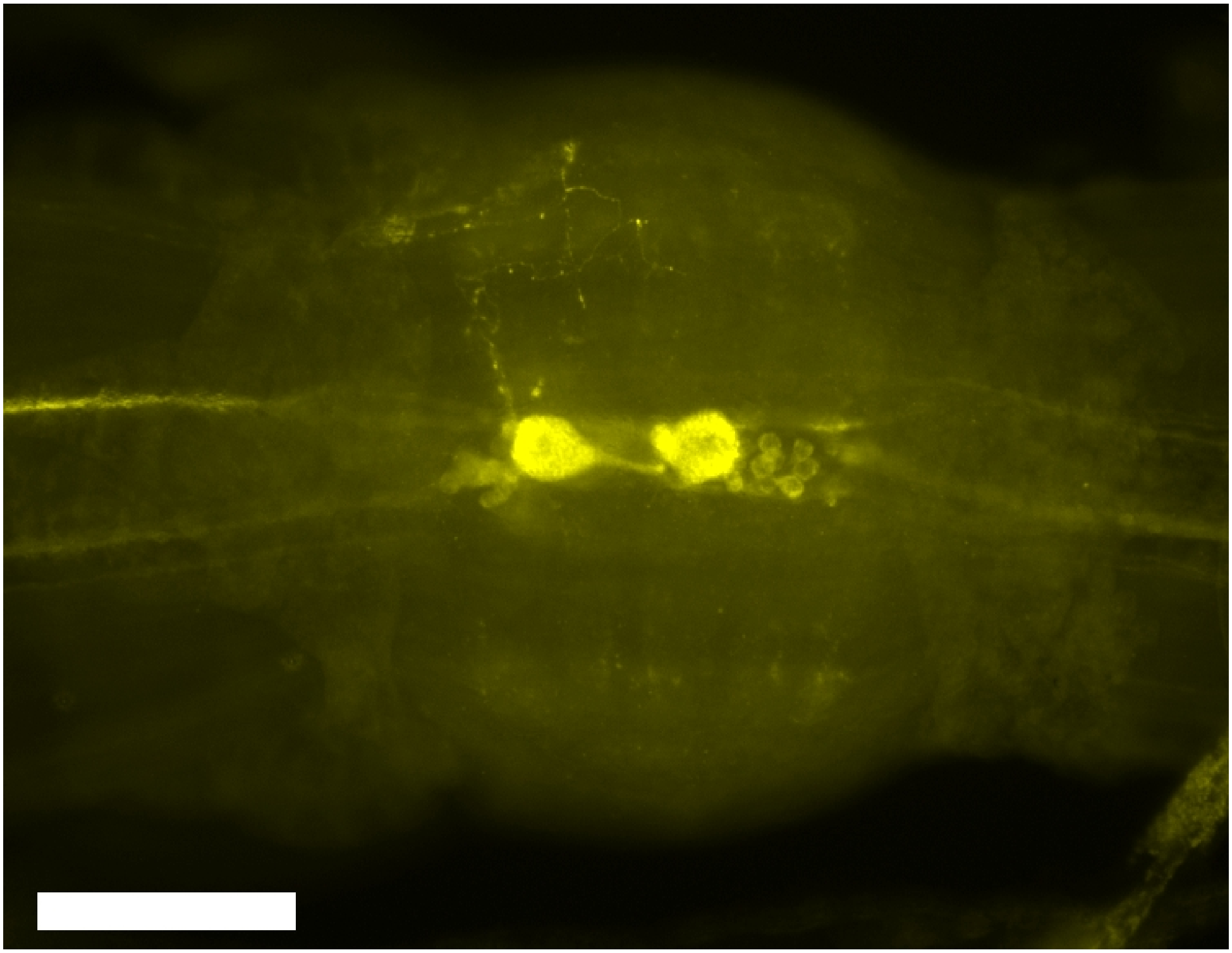

**Fig. S7.** Rabbit calcitonin A immunoreactivity on abdominal ganglion 4 from *Periplaneta americana.* Scale bar 200 µm.

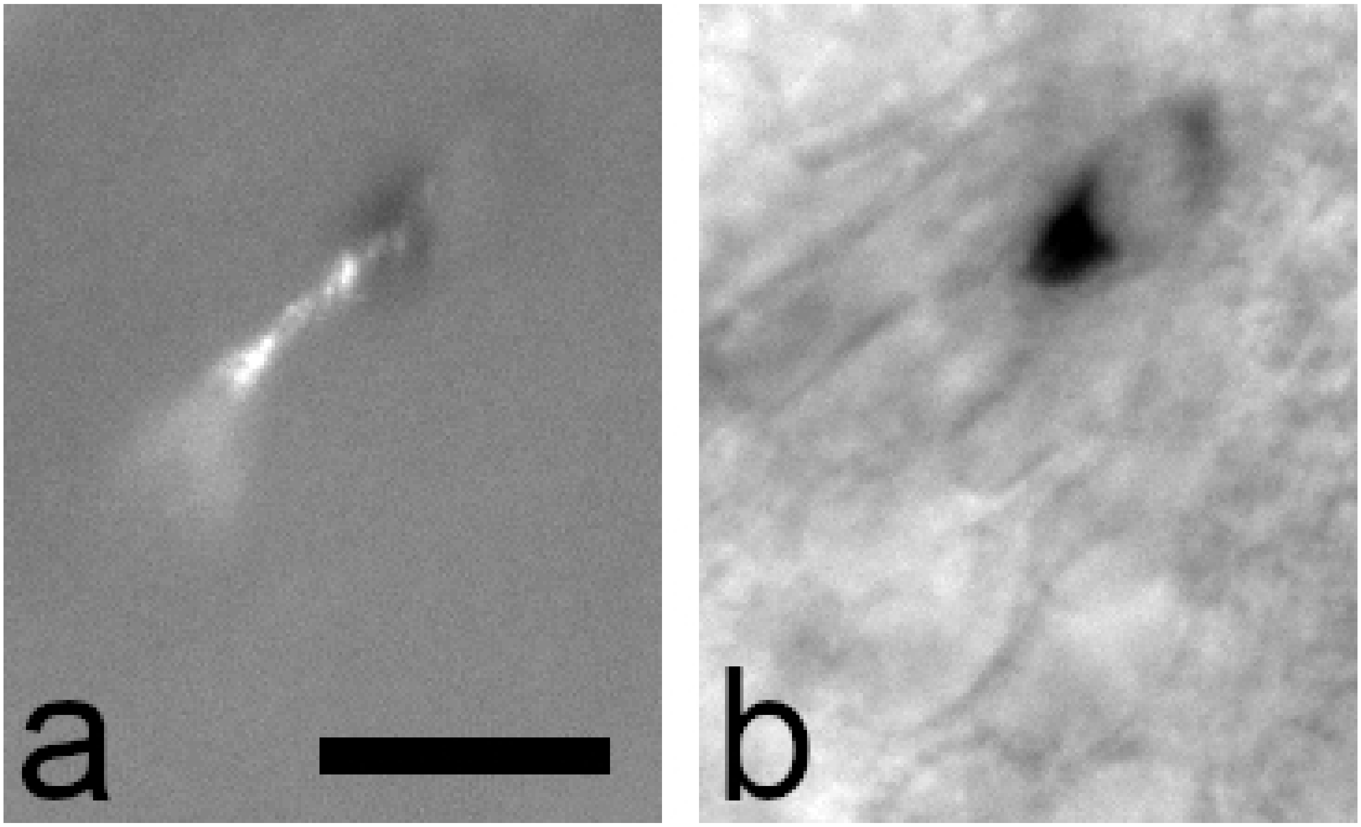


**Figure S8.** Colocalization of calcitonin B immunoreactivity in the midgut with anti-sense calcitonin B probe. Given the size of the enteroendocrine cells and the very small amount of cytoplasm double labeling of these cells is difficult to document. Here is one such cell where it is labeled with both the antiserum (a) and the anti-sense probe (b). Scale bar 25 µm.

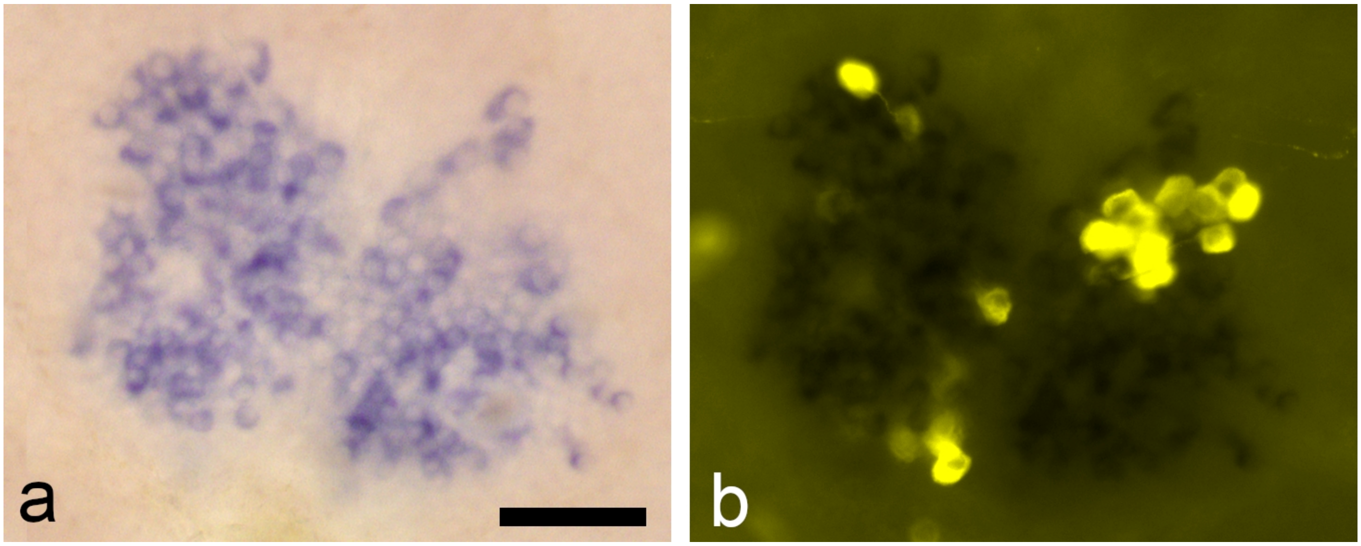


**Fig. S9.** Wholemount of *Locusta* brain showing simultaneous localization of anti-sense insulin *in situ* probe (a) and ACP immunoreactivity (b). The alkaline phosphate reaction product that is purple in (a) can be recognized as black in (b). Note the ACP cells are much larger. Scale bar 100 µm.

```
Locusta         -------MIARLL---LTLTVTAWCCYLVASQVTFSRDWSPGKRSPEPACA-----KHAATI  47
Schistocerca    -------MVARLF---LALTVTAWCCYLVTSQVTFSRDWSPGKRSPEPTCA-----KHAASI  47
Gryllus         MASWGRPLGRALLCGAAVLLVLAC---AARAQITFSRDWNAGKRAGPAPGPAPAPPL-ADHL  58
Periplaneta     -------MVHRALCWLLFLAVLSCLHPRALAQVTFSRDWNAGKRSPPPDMQCGAALKAVDQI  55


Locusta         CQLLLNELRQLAACEVKSLLRYHA----------EEVNVPVPQEIYIDGNGGR--------   90
Schistocerca    CQI-LNELRQLAACEMKSLLRYHA----------EEVNV--PQEIYIDGNGGR--------   87
Gryllus         -KSGKWPLVNSAVCSITHLTKTQIHLISSWTYTDEDKDFLLKSALHFRRNYSYKNSLSKLT  115
Periplaneta     CKVLVDEFRQLAVCETKSLLRFQR----------EIDNK--QAEIFLEGQEGR--------   96
```

**Fig. S10.** Sequence comparison of ACP precursors from *Locusta migratoria, Schistocerce gregaria, Gryllus bimaculatus* and *Periplaneta americana.* Note that the predicted mature locust ACP sequences are identical and very similar to those from *Gryllus* and *Periplaneta*, but that the latter two have a C-terminal NA-amide in stead of an SP-amide, thus explaining why these peptides are not recognized by the *Locusta* calcitonin antisera. Yellow highlighting indicates the signal peptide, red amino acid residues are those that serve as substrate for convertase and carboxypeptidase, purple residues are glycine that are predicted to be transformed in C-terminal amides and orange ones cysteines that are predicted to form disulfide bridges. Blue residues correspond to the mature ACP sequences, dark blue are those residues that are identical to the *Locusta* and *Schistocerca* sequence.
